# Invigorating human MSCs for transplantation therapy via Nrf2/DKK1 co-stimulation in a mice acute-on-chronic liver failure model

**DOI:** 10.1101/2022.05.29.493908

**Authors:** Feng Chen, Zhaodi Che, Yingxia Liu, Pingping Luo, Lu Xiao, Yali Song, Cunchuan Wang, Zhiyong Dong, Mianhuan Li, George L. Tipoe, Dongqing Wu, Min Yang, Yi Lv, Fei Wang, Hua Wang, Jia Xiao

## Abstract

Boosting stem cell resilience against an extrinsically harsh recipient environment is critical to therapeutic efficiency of stem cell-based transplantation innovations in liver disease contexts. We aimed to establish the efficacy of a transient plasmid-based preconditioning strategy to boost mesenchymal stromal cells (MSCs) capacity for anti-inflammation/antioxidant defense and paracrine actions on recipient hepatocytes. In MSCs, the master antioxidant regulator Nrf2 was found to bind directly to the antioxidant response element in the DKK1 promoter region. Activation of Nrf2 and DKK1 enhanced the anti-stress capacities of MSCs *in vitro*. In an acute-on-chronic liver failure (ACLF) murine model, transient co-overexpression of Nrf2 and DKK1 via plasmid transfection markedly improved MSC resilience against inflammatory and oxidative assaults, boosted MSC transplantation efficacy and promoted recipient liver regeneration because of a shift from the activation of the anti-regenerative IFN-γ/STAT1 pathway to the pro-regenerative IL-6/STAT3 pathway in the liver. Moreover, specific ablation of DKK1 receptor CKAP4 but not LRP6 in recipient hepatocytes nullified therapeutic benefits from MSC transplantation. In long-term observations, tumorigenicity was undetected in mice following transplantation of such transiently preconditioned MCSs. In conclusion, co-stimulation of Nrf2/DKK1 signaling decisively and safely improves the efficacy of human MSC-based therapies in mouse ACLF models through apparently CKAP4-dependent paracrine mechanisms.

## INTRODUCTION

As a vital organ governing organismal homeostasis, the liver is continually exposed to an extraordinary array of biological/chemical toxins, some of which bear profound implications for severe liver diseases. To illustrate, bacterial endotoxins (e.g. lipopolysaccharide, LPS) arising from dysregulated gut microbiota are known to drive severe hepatitis or liver failure through the portal vein (Wiest et al., 2017). Acute-on-chronic liver failure (ACLF) is a clinical syndrome defined by liver failure with pre-existing chronic liver injury. It is characterized by an acute liver insult and a rapid deterioration of liver functions and with high short-term mortality (Arroyo et al., 2016; Hernaez et al., 2017). The main etiologies for the acute liver insult include alcohol drinking, viral hepatitis (e.g. HBV), and drug-induced liver injury (DILI), while the most frequently documented etiologies for the pre-existing chronic liver injury of ACLF include chronic alcoholic consumption and HBV infection (Gustot and Jalan, 2019; Sarin and Choudhury, 2016; Zaccherini et al., 2021). Unfortunately, there is no specific effective treatment available for ACLF patients and currently treatment is based on organ support and complication resolution. Since the prognosis of ACLF probably depends on the control of bacterial infection and the recovery of multi-organ injury, early identification and treatment of the precipitating factors (e.g. bacterial infections, gastrointestinal bleeding, alcoholism, drug toxicity, and HBV reactivation) are essential (Hernaez *et al*., 2017; Sarin et al., 2019). Liver transplantation is likely the only curative treatment for ACLF patients with organ failure development in the presence of cirrhosis, especially extrahepatic organ failures (Trebicka et al., 2020). However, due to the lack of donor organs, liver transplantation is often considered to be contraindicated when the survival rate after transplantation is lower than that without transplantation (Cullaro et al., 2020). In this context, an artificial liver assist device would be desirable for spontaneous liver regeneration support and proper liver transplantation preparation. Current data showed that only therapeutic plasma exchange could improve the survival of patients with acute liver failure (and possibly ACLF). Molecular adsorbent recirculating system failed to show improvement in survival in patients with ALF or ACLF (Larsen, 2019). Therefore, the development of sustainable, cost-effective complementary treatment modalities for ACLF is urgently needed.

In recent years, exploration of human mesenchymal stromal cell (MSC) transplantation has been incentivized to treat ALF and ACLF in both experimental models and clinical patients (Kong et al., 2020; Liang et al., 2018; Shi et al., 2017; Shi et al., 2012). While still in its formative stage, MSC-based treatments have yielded generally encouraging results wherein MSCs improved recipient liver conditions by boosting hepatic regenerative capacity and exerting immuno-regulatory effects via paracrine actions on local hepatocytes (Wang et al., 2021; Yuan et al., 2019). A major challenge in achieving efficacious MSC-based therapies in the clinic lies in the poor survival rates of stem cells after transplantation (Burst et al., 2010). This problem is at least partially attributable to a pro-inflammatory and pro-oxidant host environment prevailing at the injured sites. Indeed, enhancement of endogenous antioxidant capacities of transplanted MCSs has been variously proposed to improve transplantation efficacy (Dernbach et al., 2004; Drowley et al., 2010; Zeng et al., 2015), though clinically relevant strategies remain to be established. For example, several small-molecule antioxidants or ROS (reactive oxygen species) scavengers including edaravone and N-acetyl-L-cysteine (NAC) have been explored to enhance stem cell anti-stress responses, but there has not been definite evidence that these compounds can sustainably boost MSC resilience due to a paucity of mechanistic details. Since Nrf2 (nuclear factor erythropoietin-derived 2-like 2) displays protective impact on stem cell biology in response to various environmental cues, via the regulation of pluripotency factors, redox homeostasis, aging, and cellular stress responses (Dai et al., 2020), activation of Nrf2 seems to be a plausible method for the enhancement of efficacy of stem cell transplantation (Malik et al., 2013). Unfortunately, the underlying mechanisms and subsequent profile of SC secretory factors are poorly understood. In addition, a lack of information on the direct molecular targets of transplanted MSCs’ paracrine actions also impedes advances in therapeutic inventions. While several kinds of such mechanisms have been brought to light in animal disease models, the key potential targets mediating MCS benefits call for rigorous scrutiny in molecular and biochemical approaches (Kusuma et al., 2017; Tachibana et al., 2017).

Inspired by these questions, we here endeavored to demonstrate that Nrf2 directly boosted cellular antioxidant responses and confer hepatoprotectant capabilities in part through the regulation of DKK1 secretion from human MSCs. Collectively, coordinated stimulation of the Nrf2/DKK1 signaling contributed to an enhanced anti-stress capacity of MSCs. Gratifyingly, activation of Nrf2 and DKK1 signaling via transient plasmid preconditioning of transplanted MCSs effectively and safely boosted the transplantation efficacy *in vivo* in a novel murine model of ACLF, partly due to accelerated liver regeneration because of a shift from the activation of the anti-regenerative IFN-γ/STAT1 pathway to the pro-regenerative IL-6/STAT3 pathway. In a paracrine manner, secreted DKK1 from transplanted MSC exerted pro-resolving and reparative effects on recipient mouse hepatocytes via the DKK1 receptor, cell surface cytoskeleton-associated protein 4 (CKAP4).

## RESULTS

### Nrf2 promotes the anti-stress capacity of MSC via direct regulation of DKK1

Since the transplantation of MSC preconditioned with the minocycline or doxycycline protects against ischemic injury in murine models via Nrf2 activation (Malik *et al*., 2013; Sakata et al., 2012), we first examined whether treatment of TNF-α/H_2_O_2_ (tumor necrosis factor-alpha pluses hydrogen peroxide), a well-characterized reactive oxygen intermediates- and inflammation inducer-challenged ACLF-like cell model (Bátkai et al., 2007; Gilston et al., 2001; Kudo et al., 2009), altered the basal Nrf2 activity and the release of key soluble cytokines/chemokines. A 24-h treatment with TNF-α/H_2_O_2_ significantly promoted the activity of Nrf2 in MSCs, as well as the translational and secreted levels of Wnt canonical pathway inhibitor Dickkopf-1 (DKK1) (Figure 1A). TNF-α/H_2_O_2_ treatment also enhanced the secretion of pro-inflammatory cytokines/chemokines (IL-1β, IL-6, MCP-1, and RANTES) and anti-inflammatory cytokine (IL-10) from MSCs, indicating an ACLF-comparable inflammatory environment in the cell culture system (Figure supplement 2). Next, we sought to determine whether the effects observed were because of a direct association between Nrf2 and *Dkk1* gene promoter. By using bioinformatics analysis, we found the presence of antioxidant response element sequences (AREs) at position −96 from the *Dkk1* transcription start site, which had similarity to the ARE consensus sequence observed in other Nrf2 target genes (e.g. NQO1, HMOX1, and SOD1). In consistent with the result, Nrf2-containing plasmid significantly reduced the luciferase activity of DKK1-bearing plasmid, which was hampered when DKK1 was mutated from TGACTCTGC to ATCGAGATA (Figure 1B). Moreover, a dose-dependent increase in *Dkk1* promoter activity was seen when *Nrf2* was knocked-down by siRNA (Figure 1C). Then the basal expression of *Nrf2* and *Dkk1* was overexpressed or inhibited by transfection with gene ORF (open reading frame)-bearing plasmid or shRNA, respectively, at the time of 48-h before the treatment of TNF-α/H_2_O_2_ (Figure 1D). Silencing of Nrf2 reduced cell viability in TNF-α/H_2_O_2_-challenged MSCs, whereas overexpression of Nrf2 or DKK1 evidently improved cell viability. Inhibition of DKK1 did not influence the cell viability after TNF-α/H_2_O_2_ challenge (Figure 1E). We observed consistent changes of MSCs apoptotic ratio after the manipulations of Nrf2 and DKK1 basal expression (Figure 1E). In addition, protein expression changes of the cell cycle regulator PCNA (proliferating cell nuclear antigen) and apoptotic negative regulator Bcl-2 reflected influences of Nrf2/DKK1 signaling in MSCs viability and apoptosis, respectively (Figure 1F). This result was further strengthened by the activity changes of caspase-3/8 of MSCs (Figure 1G). Since endogenous production of excessive cellular and mitochondrial reactive oxygen species (ROS) in MSCs can arise as a direct consequence of extrinsically imposed TNF-α/H_2_O_2_ toxicity (Gilston *et al*., 2001), we ventured to verify the effects of Nrf2/DKK1 modulation on oxidant-induced cellular and mitochondrial dysfunction by detection of oxidative events with CellRox (cellular oxidative stress probe) and MitoSOX (mitochondrial superoxide probe), respectively. As anticipated, exposure of TNF-α/H_2_O_2_ elevated endogenous production of cellular/mitochondrial ROS of MSCs, which was alleviated by Nrf2 or DKK1 overexpression. Knockdown of Nrf2 or DKK1 slightly exacerbated cellular/mitochondrial ROS production of MSCs (Figure 1H and 1I). Collectively, Nrf2 promotes the anti-stress capacity of MSC via direct regulation of DKK1.

**Figure 1.**
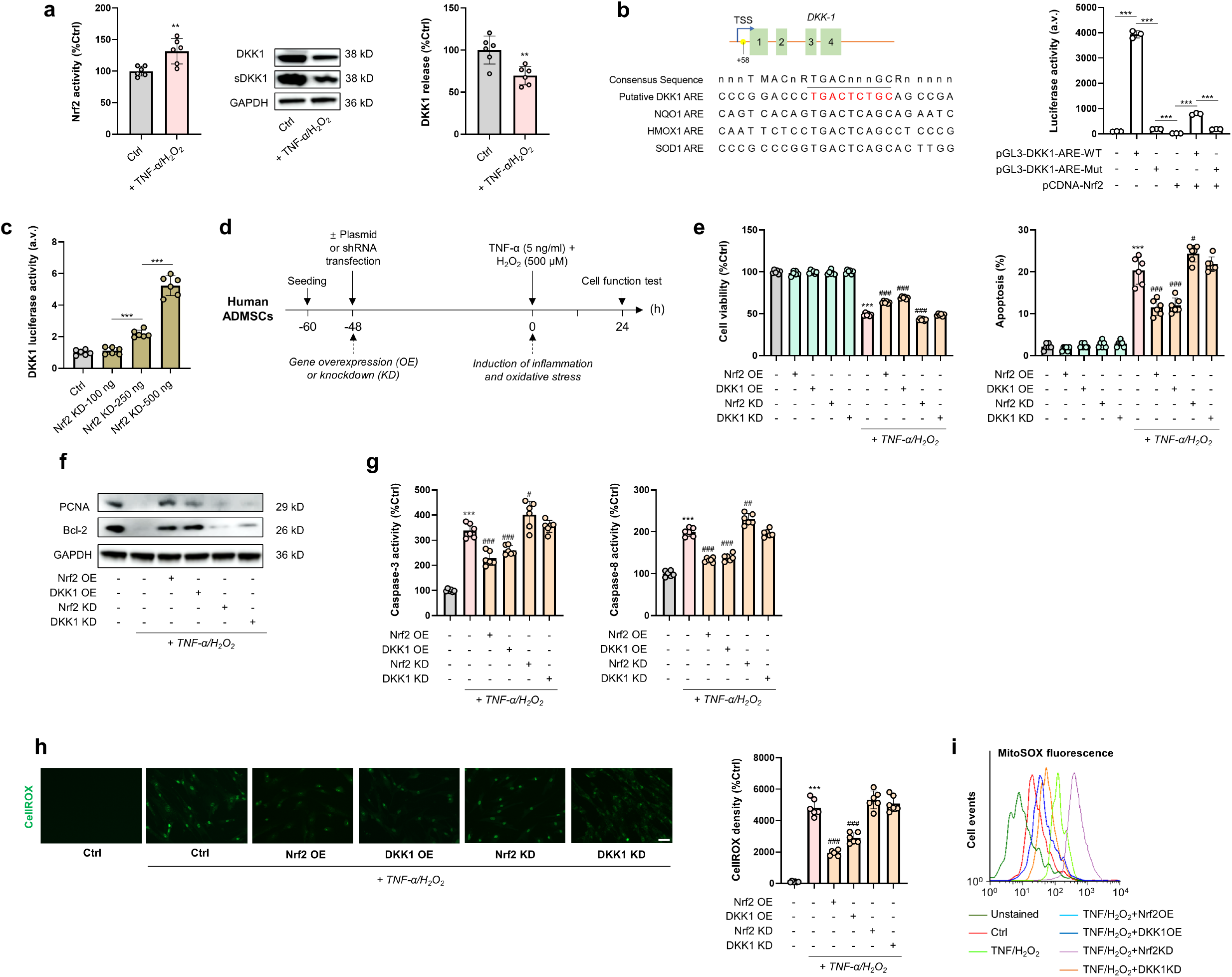
Co-stimulation of MAPK/Nrf2 and Nanog/DKK1 signaling boosts anti-stress capacity of human MSCs. (A) Left: Changes of Nrf2 activity in MSCs with or without TNF−*α*/H_2_O_2_ co-treatment. Right: Representative immunoblot results for DKK1 and secreted DKK1 (sDKK1) in MSCs with a similar test design as in panel A (*n* = 6). (B) Left: Bioinformatic analysis identified a putative ARE (antioxidant response element) located in the DKK1 promoter between −96 to −88 bp of the transcription start site (TSS). Green boxes indicate DKK1 exons. The consensus sequence for the extended ARE is shown, with the commonly identified core ARE indicated by the underlined sequence. Abbreviations used follow standard IUPAC nomenclature (M = A or C; R = A or G; Y = C or T; n = any nucleotide). Right: Relative luciferase activity of DKK1-ARE-WT and DKK1-ARE-Mut (mutant) reporter vectors in the HEK-293T cells transfected with pcDNA3.1-Nrf2 plasmid (*n* = 3). (C) Changes of DKK1 luciferase activity when Nrf2 was knocked-down by siRNA with indicated concentrations (*n* = 6). (D) Experimental design illustration showing that Nrf2 and DKK1 was overexpressed (OE) or knocked-down (KD) by transfection with gene open reading frame-bearing plasmid or shRNA, respectively, at the time of 48-h before the treatment of TNF-α/H_2_O_2_ in MSCs. (E) Changes in cell viability and apoptotic ratios of MSCs following challenge with TNF−*α*/H_2_O_2_ in the presence or absence of Nrf2/DKK1 manipulations (*n* = 6). (F) Representative immunoblot results for PCNA and Bcl-2 from MSCs following the aforementioned treatments. (G) Changes in cellular caspase-3/8 activity of MSCs with the aforementioned treatments (*n* = 6). (H) Left: Fluorescence micrographs for the detection of cellular ROS by CellROX Green and corresponding quantified fluorescence intensities in MSCs treated with TNF−*α*/H_2_O_2_ with the aforementioned treatments (*n* = 6). Scale bar = 50 μM. (i) Changes in mitochondrial superoxide levels in MSCs measured by MitoSOX in flow cytometry, following the aforementioned treatments. hADMSCs, human adipose-derived mesenchymal stromal cells. Values are expressed as mean ± SD. *, **, *** indicate *p* < 0.05, 0.01, 0.001 against an untreated MSC group (or between indicated groups), respectively; ^#, ##, ###^ indicate *p* < 0.05, 0.01, 0.001 against an TNF−*α*/H_2_O_2_ group, respectively.

### Mobilization of the Nrf2/DKK1 signaling in MSC led to a moderation of cellular/mitochondrial ROS production

In order to substantiate mechanistically how Nrf2/DKK1 signaling contributes to MSCs resilience against TNF-α/H_2_O_2_-induced stress, we validated the interrelations between this pathway and ROS production in human MSCs by using MitoQ (mitochondrial ROS scavenger) and N-acetylcysteine (NAC; total cellular antioxidant) in the presence of TNF-α/H_2_O_2_. Consistent with our assumptions, co-treatment with MitoQ or NAC significantly ameliorated TNF-α/H_2_O_2_-induced cell damages in several aspects, including an increase in cell viability/PCNA expression and a reduction in apoptosis and cellular/mitochondrial ROS production (Figure 2A-2C). Nrf2 activity of MSCs was increased by TNF-α/H_2_O_2_ exposure, but was re-balanced by co-treatment with MitoQ or NAC. Similarly, TNF-α/H_2_O_2_ challenge evidently reduced p38 MAPK phosphorylation and DKK1 protein expression, which were substantially restored by MitoQ/NAC co-treatment (Figure 2D). It is noteworthy that the decreased ERK phosphorylation seen during TNF-α/H_2_O_2_ exposure was further suppressed by MitoQ/NAC (Figure 2D). Importantly, previous studies have postulated cross-regulation between MAPKs and DKK1 in cancer cells and T cells (Browne et al., 2016; Chae et al., 2016; Rachner et al., 2015), but it remains largely unknown whether potential crosstalk exists between the MAPK and Nrf2/DKK1 pathways in human MSCs. In our *in vitro* study on TNF-α/H_2_O_2_-induced cell damages, we found that the provoked Nrf2 activity were further enhanced by SB203580 (p38 MAPK inhibitor) or UO126 (MEK1/2 inhibitor). Change of cellular and secreted DKK1 levels was in opposite to that of Nrf2, further confirmed the negative regulatory loop between Nrf2 and DKK1. When Nrf2 was overexpressed by plasmid transfection, both basal and TNF-α/H_2_O_2_-suppressed phosphorylated p38 MAPK levels were markedly elevated. Contrarily, Nrf2 overexpression strongly suppressed ERK phosphorylation under basal and TNF-α/H_2_O_2_-treated conditions (Figure 2E). Overexpression of DKK1 in MSCs visibly increased phosphorylated p38 MAPK levels but decreased phosphorylated ERK levels with or without TNF-α/H_2_O_2_ exposure (Figure 2E). Moreover, TNF-α/H_2_O_2_ exposure diminished the cellular levels of NAD(P)H dehydrogenase [quinone] 1 (NQO-1) and heme oxygenase-1 (HO-1) (Figure supplement 3A), which are Nrf2-regulated antioxidant enzymes important to stem cell homeostasis (Chen et al., 2014). Pharmacological inhibition of p38 MAPK and ERK further reduced and restored their expression, respectively, while Nrf2 overexpression significantly improved both the basal and TNF-α/H_2_O_2_-repressed levels of NQO-1 and HO-1 (Figure supplement 3A). When endogenous expression of NQO-1 or HO-1 in either type of human MSCs was silenced by specific shRNAs, the ameliorative effects of Nrf2 or DKK1 overexpression on cell injury were drastically curtailed (Figure supplement 3B-3C), suggesting a dependence on NQO-1 and HO-1 as distal mediators in MSC anti-stress response. Collectively, mobilization of the Nrf2/DKK1 signaling in MSC led to a moderation of cellular/mitochondrial ROS production, partly via the antioxidant actions of NQO-1 and HO-1.

**Figure 2.**
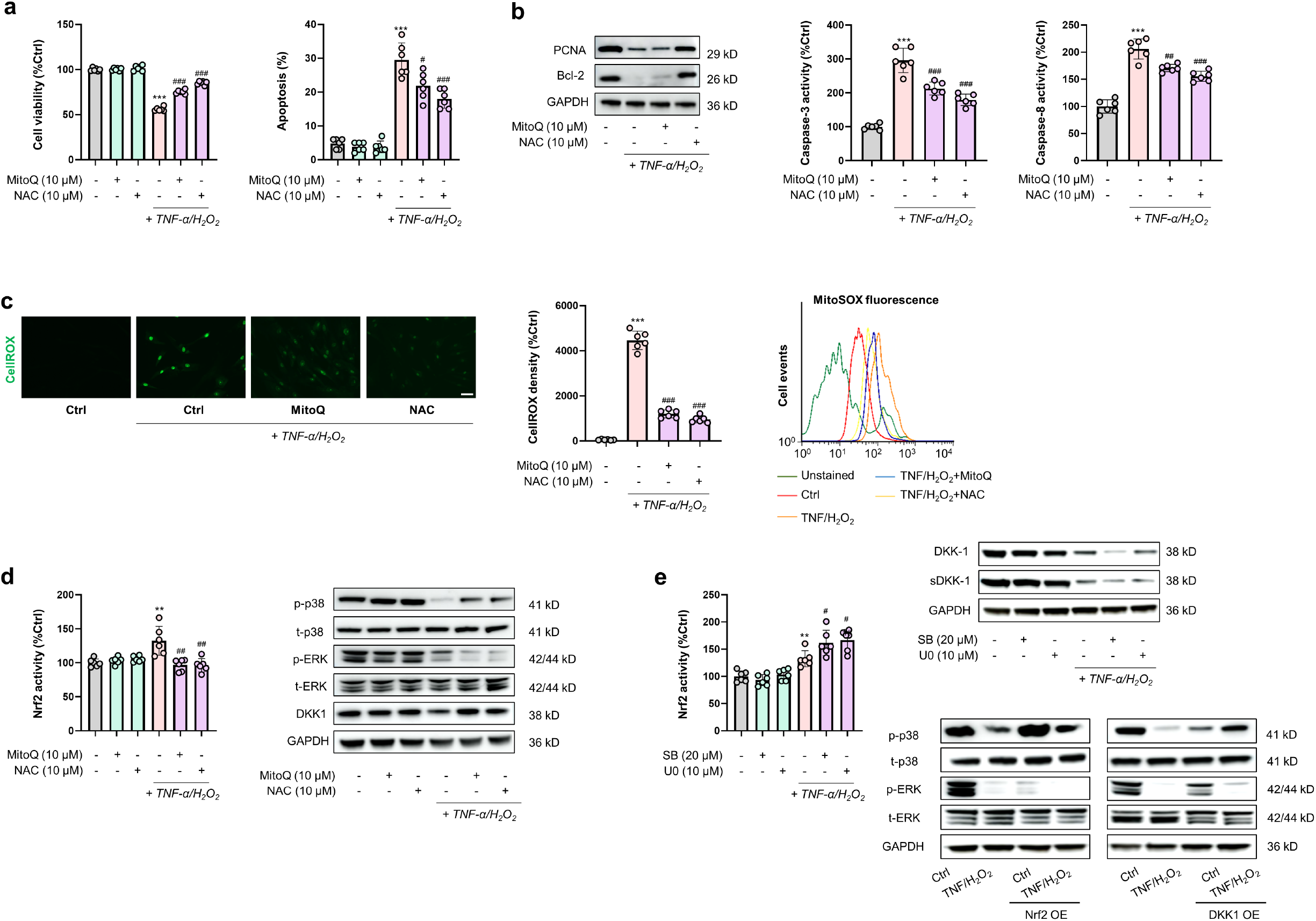
Enhanced MSCs resilience through co-stimulation of MAPK/Nrf2 and DKK1 signaling is attained by reducing ROS generation. (A) Cell viability and apoptotic ratios changes of MSCs treated with TNF−*α*/H_2_O_2_ in the presence or absence of MitoQ or NAC (*n* = 6). (B) Representative immunoblotting results for PCNA and Bcl-2 in MSCs (left) and changes in cellular caspase-3/8 activity of MSCs with a similar test design as in panel A *(n* = 6). (C) (Left) Fluorescence micrographs for the detection of cellular ROS by CellROX Green and corresponding quantified fluorescence intensities in MSCs following antioxidant intervention (*n* = 6) (Scale bar = 50 μM). (Right) Changes in mitochondrial superoxide levels in MSCs following antioxidant intervention, as measured by MitoSOX in flow cytometry (*n* = 6). (D) Assays on transcriptional activities of Nrf2 (left), and representative immunoblot results (right) for phosphorylated p38 MAPK (p-p38), total p38 MAPK (t-p38), p-ERK, t-ERK, and DKK1 in MSCs following antioxidant intervention (*n* = 6). (E) (Left) Changes in transcriptional activities of Nrf2 in MSCs following TNF−*α*/H_2_O_2_ challenge in the presence or absence of MAPK/ERK inhibitors (*n* = 6). (Right-upper) Representative immunoblot results for DKK1 and secreted DKK1 (sDKK1) in MSCs following the aforementioned treatments. (Right-lower) Representative immunoblot results for p-p38, t-p38, p-ERK, t-ERK of MSCs following TNF−*α*/H_2_O_2_ challenge, with or without Nrf2/DKK1 overexpression (OE). Data are expressed as mean ± SD. **, *** indicate *p* < 0.01 and *p* < 0.001 against an untreated MSC group, respectively; ^#, ##, ###^ indicate *p* < 0.05, 0.01, 0.001 against a TNF−*α*/H_2_O_2_ group, respectively. SB, SB203580, the inhibitor of p38 MAPK; U0, U0126, the inhibitor of ERK.

Transient co-overexpression of Nrf2 and DKK1 boosts MSC resistance against stress Although Nrf2 directly and negatively regulates DKK1 expression, and *vice versa*, we found overexpression of each of them could alleviated inflammation- and oxidative stress-induced cell injury in MSCs. To maximize the alleviative effects and to avoid the negative regulating loop between them, we attempted to co-overexpress Nrf2 and DKK1 by constructing an optimized pIRES2-Nrf2-DKK1 expression plasmid (Figure 3A) and transfected MSCs with it for 5 days (approximate the same duration where MSCs need to endure the harsh effects of local stress following transplantation) to verify its protein inductive potential. Following transfection, protein expression of Nrf2, total DKK1 and secreted DKK1 in human MSCs began to rise at day 2, peaked at day 3 and returned to near-basal levels at day 5 post-transfection (Figure 3B). Importantly, transient transfection of this plasmid did not alter MSC viability or MSC differentiation potential (Figure supplement 4). Based on this observation, we subjected MSCs to added TNF-α/H_2_O_2_ challenge at day 3 post-transfection for another 24 h to evaluate the cells’ anti-stress capacity gained from plasmid transfection. As anticipated, pIRES2-Nrf2-DKK1 plasmid transfection significantly restored cell viability, reduced apoptosis, and dampened cellular/mitochondrial ROS production in MSCs without altering their basal status (Figure 3C-3E).

**Figure 3.**
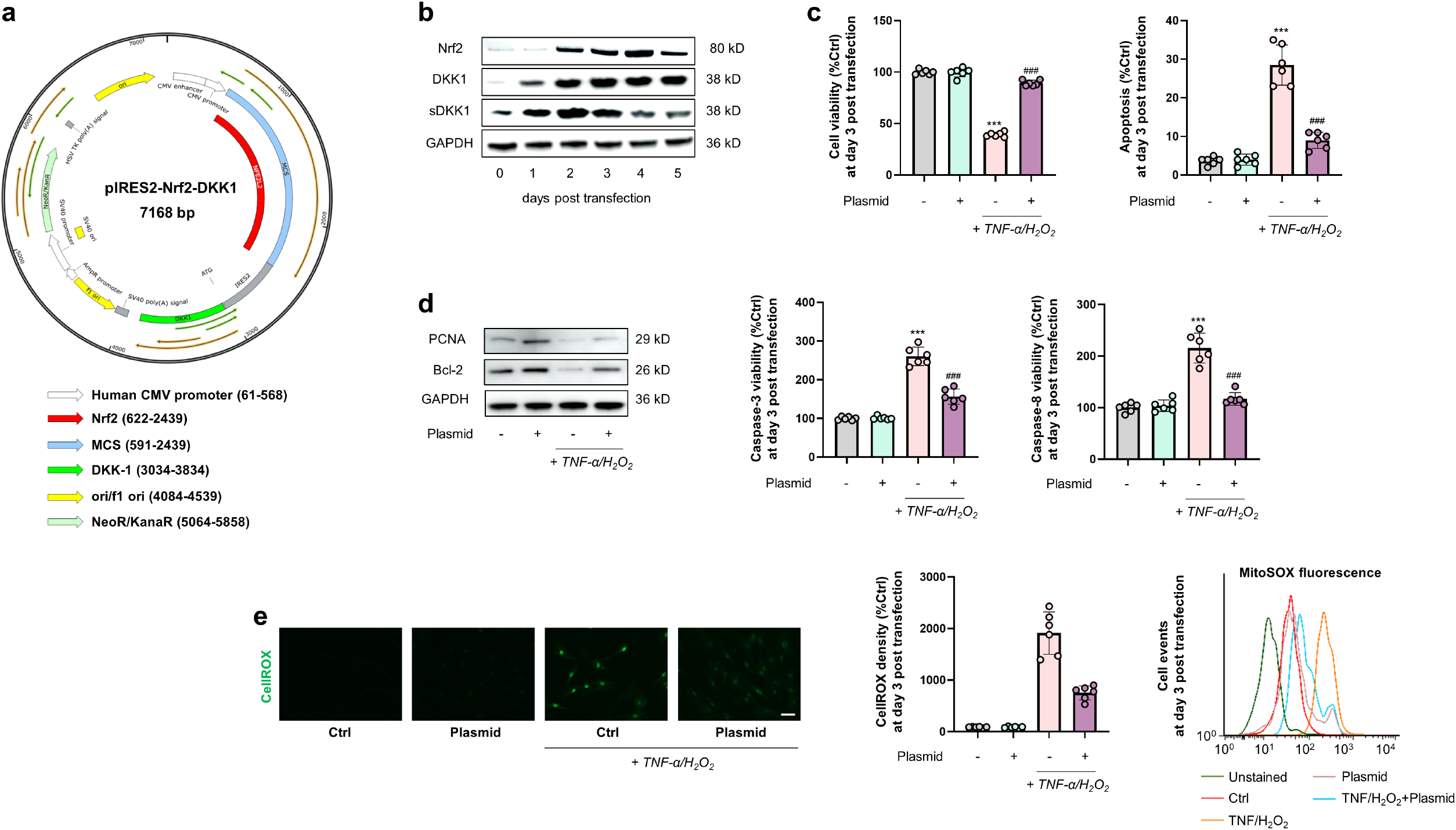
Preconditioning by co-overexpression of Nrf2 and DKK1 enhances MSC resistance to exogenous stress. (A) Plasmid map for the constructed pIRES2-Nrf2-DKK1. (B) Representative immunoblot results for time-lapse study (day 0 - day 5) on Nrf2, DKK1 and secreted DKK1 (sDKK1) expression in MSCs following transfection of the pIRES2-Nrf2-DKK1 plasmid. (C) Changes in cell viability (Left) and apoptotic ratios (Right) of MSCs following TNF−*α*/H_2_O_2_ challenge with or without transfection of the pIRES2-Nrf2-DKK1 plasmid (*n* = 6). (D) (Left) Representative immunoblotting results for PCNA and Bcl-2 in MSCs and (Right) changes in cellular caspase-3/8 activity of MSCs following TNF−*α*/H_2_O_2_ challenge with or without transfection of the pIRES2-Nrf2-DKK1 plasmid *(n* = 6). (E) (Left) Detection of cellular ROS in MSCs by CellROX Green and corresponding quantified fluorescence intensities in MSCs following TNF−*α*/H_2_O_2_ challenge with or without transfection of the pIRES2-Nrf2-DKK1 plasmid (*n* = 6) (Scale bar = 50 μM). (Right) Changes in mitochondrial superoxide levels in MSCs, following TNF−*α*/H_2_O_2_ challenge with or without transfection of the pIRES2-Nrf2-DKK1 plasmid (*n* = 6). Data are expressed as mean ± SD. *** indicates *p* < 0.001 against an untreated MSC group; ^###^ indicates *p* < 0.001 against an TNF−*α*/H_2_O_2_ group.

### Co-stimulation of Nrf2 and DKK1 expression improves MSC transplantation efficacy in an ACLF mice model

By using a novel ACLF mice model combining chronic liver injury, acute hepatic insult, and bacterial infection that phenocopies some of the key clinical features of ACLF patients (Xiang et al., 2020), we tested the MSC-protective effects of Nrf2/DKK1 co-stimulation strategy (Figure 4A). Empirically, both death counts of ACLF mice (during a 9-d time window) were evidently mitigated by the transplantation of human MSCs. In particular, preconditioning MSCs with the pIRES2-Nrf2-DKK1 plasmid prior to transplantation further enhanced the protective effects in either type of mice. No further mouse death was seen in all groups during an extended 9-day observation period (Figure 4B). High levels of serum ALT and total bilirubin (TBIL), elevation of circulating neutrophils and blood urea nitrogen (BUN), and reduction of renal microvascular flow were all significantly alleviated by MSCs transplantation, and further strengthened by pIRES2-Nrf2-DKK1 plasmid preconditioning (72-h post *K.P*. administration data presented here; Figure 4C and 4D). ACLF-induced severe pathological liver damages including necrosis, inflammation, fibrosis, and lipid peroxidation were also ameliorated by MSCs and plasmid preconditioning (Figure 4E). Successfully homed human MSCs in the damaged mice liver were labelled with human nuclear antigen (hNA) staining, DKK1 protein, and alpha-1 antitrypsin (αAT) protein which showed that transfection with the pIRES2-Nrf2-DKK1 plasmid significantly improved the homing efficacy of MSCs, which was closely associated with the accelerated liver regeneration, as demonstrated by hepatic Ki-67 staining (Figure 4F). Since this ACLF model was reported to induced evident liver fibrosis, we also investigated the possible fibrotic amelioration after MSCs transplantation.

**Figure 4.**
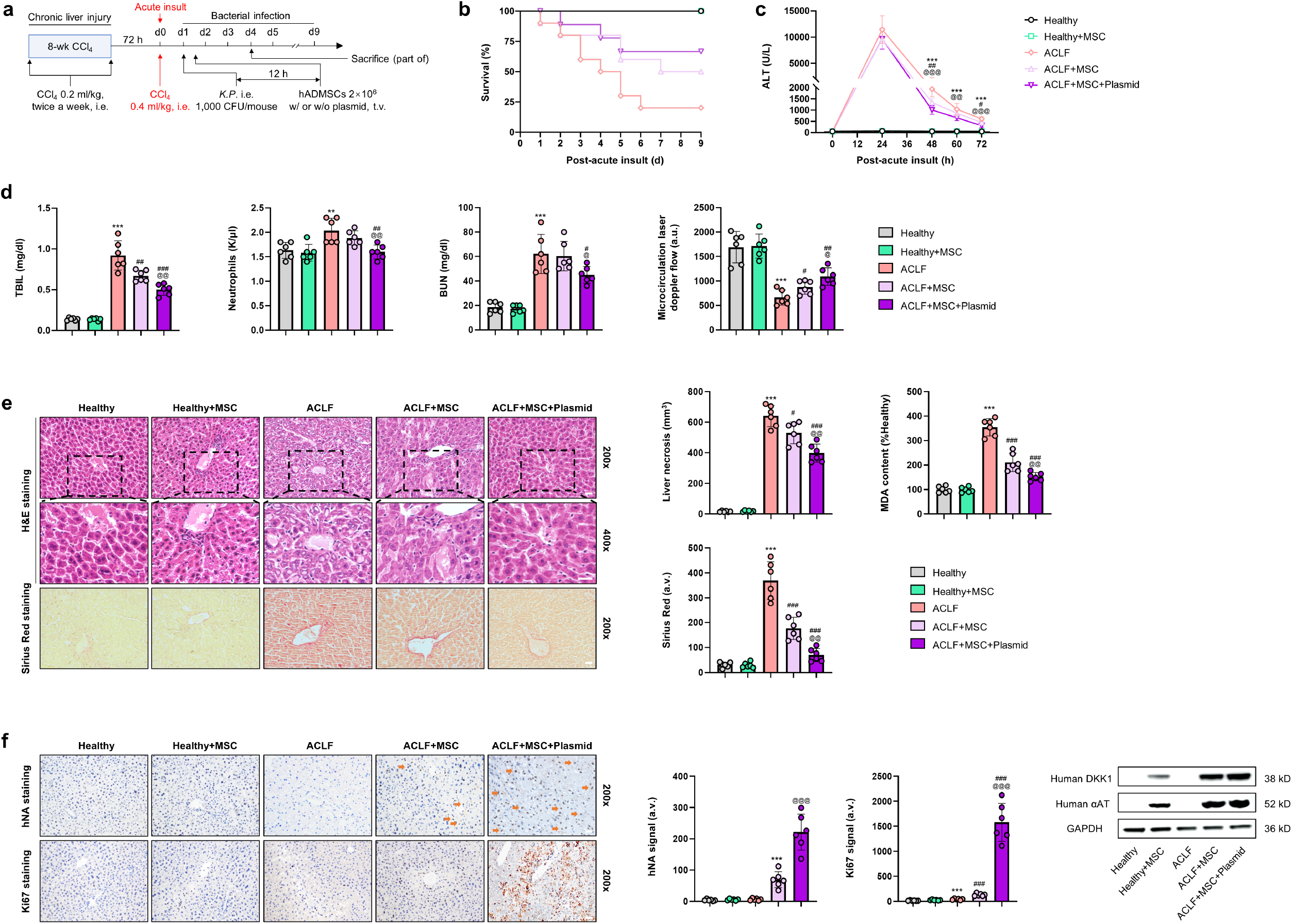
Preconditioning with a pIRES2-Nrf2-DKK1 plasmid improves transplantation efficacy of MSCs in a murine model of acute-on-chronic liver failure (ACLF). (A) Schematic timeline of the ACLF mice model establishment with carbon tetrachloride (CCl_4_) and *Klebsiella pneumoniae* (K.P.) injection with or without MSC of Nrf2/DKK1 co-stimulation. (B) Survival counts of ACLF mice with or without injection of human MSCs (naive or pre-transfected with pIRES2-Nrf2-DKK1) for 9 days (*n* = 10 per group). (C) Changes in serum ALT levels in mice as depicted in panel A (72-h post *K.P*. administration; *n* = 8). (D) Changes in liver total bilirubin (TBIL), circulating neutrophils, blood urea nitrogen (BUN), and renal microvascular flow in mice as depicted in panel A (72-h post *K.P*. administration; *n* = 6). (E) Representative images of H&E and Sirius Red staining of the mice liver, and corresponding quantification of liver necrosis areas and liver fibrosis areas, and changes in liver MDA contents in mice as depicted in panel A (72-h post *K.P*. administration; *n* = 6). (F) Representative immunohistochemical results for human nuclear antigen (hNA) and Ki67 staining and representative immunoblot results for human DKK1/αAT in the liver from mice as depicted in panel A (72-h post *K.P*. administration; *n* = 6). Scale bar = 50 μM. Arrows indicate typical IHC signals. Data are expressed as mean ± SD. **, *** indicate *p* < 0.01, 0.001 against a healthy group, respectively; ^#, ##, ###^ indicate *p* < 0.05, 0.01, 0.001 against an ACLF group, respectively; ^@, @@, @@@^ indicate *p* < 0.05, 0.01, 0.001 against an ACLF-challenged plasmid-naïve MSC group, respectively.

### MSCs preconditioned with pIRES2-Nrf2-DKK1 resolves ACLF injury by enhancing the pro-regenerative IL-6/STAT3 pathway but attenuating the anti-regenerative IFN-γ/STAT1 pathway

To investigate the mechanisms for MSCs transplantation-induced liver regeneration and fibrosis resolution, we measured serum cytokines and found that in ACLF mice, IL-6 protein level was significantly inhibited while IFN-γ protein level was significantly provoked when compared with that of control mice. In the liver, the mRNA level changes of *Il6* and *Bcl2* corresponded with that of serum IL-6 protein, while *Ifna* and *Stat1* mRNA level changes corresponded with that of serum IFN-γ protein (Figure 5A and 5B). Western blot analyses revealed that ACLF mice had enhanced phosphorylated levels of STAT1 and STAT3 vs. control mice, which was suppressed after the transplantation with MSCs, with or without pIRES2-Nrf2-DKK1 plasmid preconditioning (Figure 5C and 5D). In contrast to STAT activation, the expression of cyclin D1 was lower in mice with ACLF compared to those control mice (Figure 5C and 5D).

**Figure 5.**
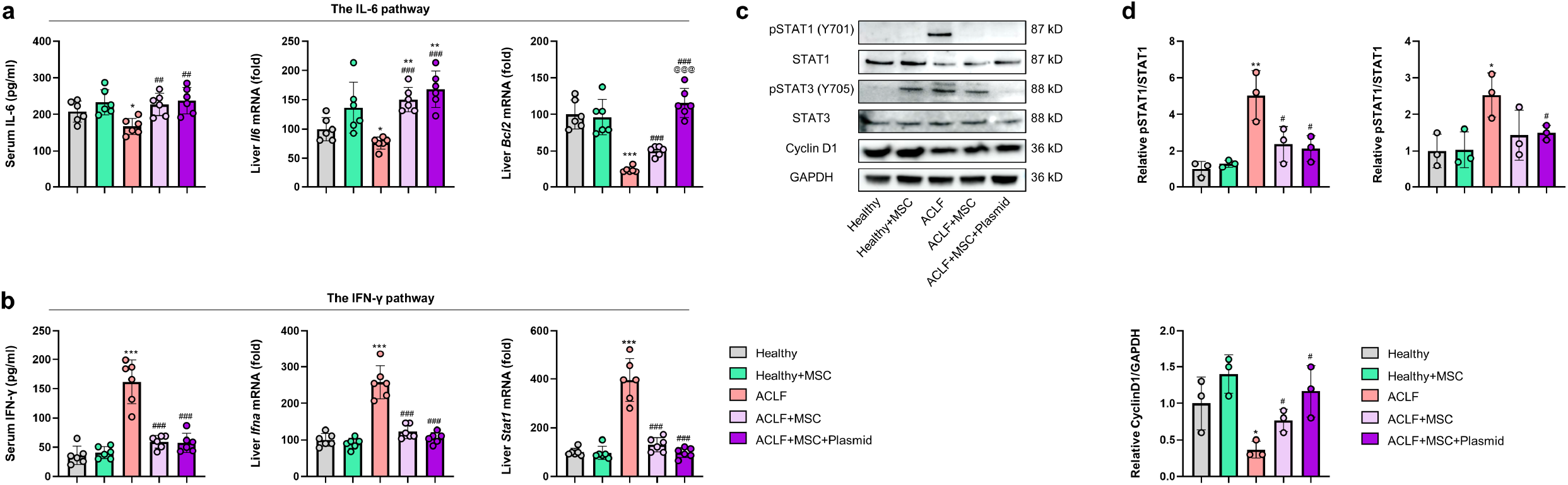
MSCs preconditioned with pIRES2-Nrf2-DKK1 resolves ACLF injury by enhancing the pro-regenerative IL-6/STAT3 pathway but attenuating the anti-regenerative IFN-γ/STAT1 pathway. (A,B) Serum IL-6 and IFN-γ, and relative mRNA expressions of *Il-6* and *Ifng*, and their downstream target genes (*Bcl2* and *Stat1*, respectively) in the ACLF or control mice after the transplantation with MSCs, with or without pIRES2-Nrf2-DKK1 plasmid preconditioning *(n* = 6). (C) Liver extracts were subjected to Western blot analysis of phosphorylated STAT1 (pSTAT1), STAT1, pSTAT3, STAT3 and Cyclin D1 in mice as depicted in panel A. (D) Relative quantification of STAT1 and STAT3 (the phosphorylated level was divided by the total protein level) and Cyclin D1 in mice as depicted in panel A *(n* = 3). Data are expressed as mean ± SD. *, **, *** indicate *p* < 0.05, 0.01, 0.001 against a healthy group, respectively; ^#, ##, ###^ indicate *p* < 0.05, 0.01, 0.001 against an ACLF group, respectively; ^@@@^ indicates *p* < 0.001 against an ACLF-challenged plasmid-naïve MSC group.

### Hepatocyte membrane receptor CKAP4 but not LRP6 mediates MSC-based recipient liver repair

As a canonical inhibitor of Wnt signaling, DKK1 has previously been shown to be secreted by MSCs to ameliorate tissue injury or reduce liver fibrosis (Prockop et al., 2003; Yang et al., 2017). In a similar vein, we speculated whether DKK1 receptor(s) on hepatocyte cell membranes transduces DKK1 signaling from transplanted human MSC to promote repair processes in murine recipient hepatocytes. Thus, by using AAV8-ligated shRNAs, we knocked down the expression two well-documented hepatocyte DKK1 receptors, cytoskeleton-associated protein 4 (CKAP4) and low-density lipoprotein receptor–related protein 6 (LRP6) (Kimura et al., 2016), specifically in the liver (Figure 6A). Compared with their WT littermates, LRP6 conditional knockdown (CKD) mice showed a similar death rate, while CKAP4 CKD mice had a greater death rate upon ACLF challenge (Figure 6B). Changes in serum ALT, liver histology, and hepatic injury markers were consistent (72-h post *K.P*. administration data presented here; Figure 6C and 6D; WT mice data not shown here). Of note, hepatic knockdown of CKAP4 or LRP6 did not influence the hepatic hNA signal, as well as human DKK1 protein and αAT protein levels, indicating that the homing efficacy of MSCs transplantation, with or without pIRES2-Nrf2-DKK1 preconditioning, did not rely on hepatocyte CKAP4 and LRP6 (Figure 6E). In contrast, hepatic staining of Ki-67, protein expression analysis on p-Akt, cyclin D1, and PCNA demonstrated that hepatic knockdown of CKAP4, when compared with LRP6 knockdown or WT mice, exhibited significantly lower level of liver regeneration, which suggests that MSC-based intervention for recipient liver regeneration was CKAP4-dependent (Figure 6F and 6G).

**Figure 6.**
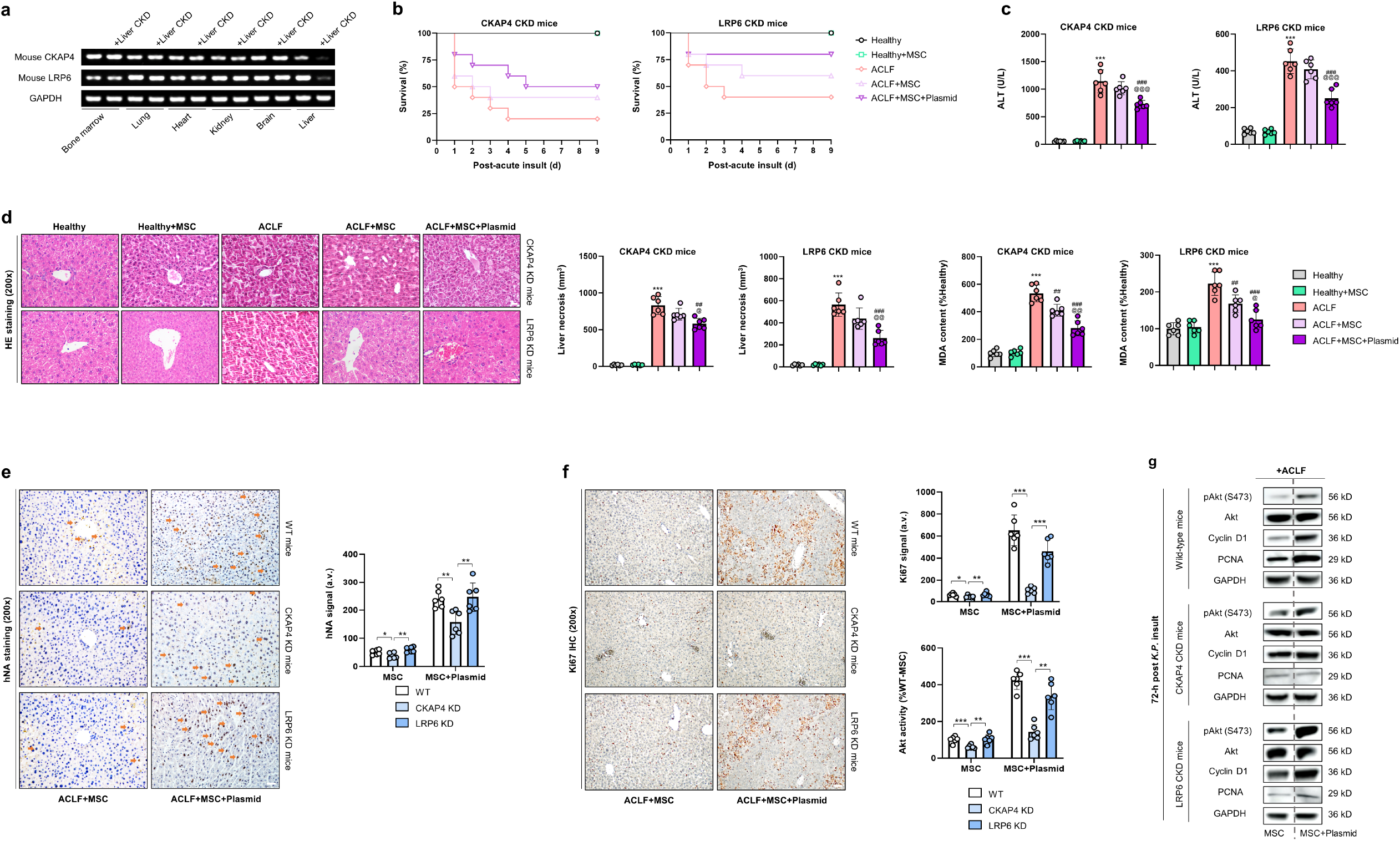
The DKK1 receptor CKAP4, but not LRP6, in host hepatocytes is a paracrine target of MSC-based therapy. (A) Representative genotyping results for hepatic-specific knockdown of CKAP4 and LRP6 by AAV8-mediated shRNA in mice (in the tissue extracts of bone marrow, lung, heart, kidneys, brain, and liver). (B) Survival counts of mice with ACLF challenge with or without injection of human MSCs (naive or pre-transfected with pIRES2-Nrf2-DKK1) for 9 days, following hepatic knockdown (CKD) of CKAP4 or LRP6 (*n* = 10 per group). (C) Changes in serum ALT levels in mice as depicted in panel B (72-h post *K.P*. administration; *n* = 6). (D) Representative images for liver H&E staining and corresponding quantification of liver necrosis areas and liver fibrosis areas, and changes in liver MDA contents in mice as depicted in panel B (Dashed lines indicate typical necrotic areas in the liver. 72-h post *K.P*. administration; *n* = 6). (E) Representative images for liver human nuclear antigen (hNA) immunohistochemical staining and corresponding quantification of hNA density in mice as depicted in panel B (72-h post *K.P*. administration; *n* = 6). Arrows indicate typical IHC signals. (F) Representative images for liver Ki67 immunohistochemical staining and corresponding quantification of Ki67 density, and enzyme-immuno assay measurements of hepatic Akt activity in mice as depicted in panel B (*n* = 6). (G) Representative immunoblot results for liver phosphorylated Akt (p-Akt), Akt, cyclin D1, and PCNA in mice with or without knockdown of hepatic CKAP4 or LRP6. Scale bar = 50 μM. Data are expressed as mean ± SD. *** indicates *p* < 0.001 against a healthy group; ^##, ###^ indicate *p* < =0.01, 0.001 against an ACLF group, respectively; ^@, @@, @@@^ indicate *p* < 0.05, 0.01, 0.001 against an ACLF-challenged plasmid-naïve MSC group, respectively. For panels E and F, *, **, *** represent *p* < 0.05, 0.01, 0.001 between indicated groups.

### Long-term transplantation with plasmid-transfect MSCs is safe in murine models

Potential risks for tumorigenicity are an important safety issue in the development of MSC-based therapies, particularly for viral- or plasmid-manipulated MSCs (Tolosa et al., 2016). In our ACLF model, no mice developed tumor (tumor incidence rate: 0%) during their long-term observation period (24 weeks). In contrast, all animals in the positive control groups of healthy and ACLF models showed severe symptoms of dyspnea and minimal activity from 5-6 weeks after ES-3D cell injection. Gross morphology suggests that 100% of the mice developed tumors in the lungs (Table supplement 1). To assess the long-term viability of donor MSCs, human albumin was measured in the mouse serum at 12- and 24-weeks post-transplantation. Human albumin levels were determined to be about 2 times higher in the serum of ACLF mice transplanted with Nrf2/DKK1 preconditioned MSCs than mice with plasmid-naïve MSCs at 12-week post-injection. By the 24^th^ week, the differences in human albumin levels between mice transplanted with Nrf2/DKK1 preconditioned MSCs and mice with plasmid-naïve MSCs became less pronounced than in the 12^th^ week (Figure supplement 5).

## Discussion

Accumulating evidence converges on the proposition that enhancement of anti-stress capacity of transplanted stem cells favorably influences therapeutic outcomes in a variety of diseases, though the essential signaling pathways that regulate mechanisms therein remain incompletely understood. Previous studies have shown that Nrf2 and its upstream MAPK pathways are involved in antioxidant-promoted stem cell resistance against exogenous stress, but their exact roles in reparative processes in liver injury require further elucidation (Drowley *et al*., 2010; Zeng *et al*., 2015). In this context, we demonstrate here that the regulatory roles of p38 MAPK and ERK in human MSC injury are opposite. p38 MAPK inhibition worsened while ERK inhibition alleviated TNF-α/H_2_O_2_-induced cell injury. Nrf2 directly bound to the ARE element of the promoter of DKK1, which was shown to be indispensable for protecting transplanted MCS against local stress. Although there was a negative regulation between Nrf2 and DKK1, overexpression of them simultaneously exhibited the best protective effects on MSCs than that of overexpression for any of them. Further analysis shows that Nrf2/DKK1 overexpression attenuated cellular/mitochondrial ROS production and restoration of antioxidant reserves. In addition, augmented expression of Nrf2 and DKK1 increased the levels of phosphorylated p38 MAPK and reduced the levels of phosphorylated ERK, to form a positive feedback loop that sustains anti-stress regulation within preconditioned MSCs. This finding is in agreement with several previous studies supporting a direct crosstalk between MAPK and Nrf2/DKK1 in other cell types (Browne *et al*., 2016; Kim et al., 2014; Naidu et al., 2009; Niwa et al., 2009).

Selective overexpression of key proteins to boost the anti-stress capacity of MSCs prior to transplantation is a theoretically sound strategy for improving therapeutic efficacy. In application contexts elsewhere, for example, forced myocardin expression in human MSCs by adenoviral gene transfer promoted their cardiomyogenic differentiation and transplantation efficiency in murine ischemic heart injury models (Grauss et al., 2008). Co-expression of the HCMV proteins US6 and US11 through retroviral vector-based transfection in human MSCs successfully switched off recognition of MSCs by the immune system, thus allowing a higher level of productive engraftment in the murine liver after transplantation (Soland et al., 2012). Nevertheless, since viral gene transfer methods genetically drive host cell reprogramming via modulation of activities of target and neighboring genes at the insertion site, they potentially raise safety issues of possible tumor development post-transplantation, especially in clinical applications (Wang et al., 2014). In comparison, transiently induced overexpression of target gene(s) in MSCs by transfection with carefully conceived plasmids seem to be a relatively safe and technically worthy method. Indeed, as compellingly demonstrated in this study, this alternative approach can significantly ameliorate therapeutic outcomes in our clinically-relevant ACLF model through the use of Nrf2/DKK1 preconditioned MSCs. Long-term observations following transplantation did not suggest any adverse effects, precluding the possibility of carcinogenesis.

In terms of mechanisms, the recipient liver signaling cascades directly molded by stem cell therapies after injury have long been an enigma. It has been proposed that stem cells are capable of orchestrating host hepatocyte regeneration via direct homing, replenishment of functional hepatocytes, and paracrine actions (e.g. via secreting proteins and extracellular vesicles for cell-to-cell communication). CKAP4 and LRP6 were reported to be the direct DKK1 receptors implicated in cancer cell proliferation, with similar affinity but distinct cysteine-rich domains (Kimura *et al*., 2016). In this current study, we asked which of the receptors matter and found that CKAP4, but not LRP6, mediated the DKK1-induced host hepatocyte regeneration partly through Akt activation in the ACLF hepatotoxicity model. Indeed, we believe that whether other DKK1 targets in hepatocytes contribute to this intricate process warrants further investigation. As an important point to note, however, abundant DDK-1 expression may promote hepatocellular carcinoma cell migration and invasion and serve as a protein biomarker of liver cancers (Chen et al., 2013; Shen et al., 2012). Thus, transient overexpression of DKK1 as an intervention strategy should be viewed with caution for some patients with such cancers. Insights gained on related signaling pathways underpinning the anti-stress capacity of MSCs will definitively help improve the efficacy and technical maturity of MSC-based transplantation therapies for other types of intractable clinical diseases.

In conclusion, we have herein demonstrated that the Nrf2/DKK1 signaling pathway sustains and enhances MSC resilience against extrinsically imposed stress post-transplantation. The MAPK member proteins p38 and ERK are also involved in this process. MSC preconditioning by transiently induced co-overexpression of Nrf2 and DKK1 via plasmids efficaciously and safely improved the transplantation efficacy and therapeutic outcomes of human MSCs in a murine ACLF model. It is hoped that our findings will help lay a theoretical foundation for further innovations of translationally mature MSC-based therapies for liver diseases.

## EXPERIMENTAL PROCEDURES

### MSC isolation, culture and validation of surface markers

Commercially available human adipose-derived mesenchymal stromal cells (hADMSCs; #HUXMD-01001) were purchased from Cyagen Biosciences (Guangzhou, China), and handled according to the manufacturer’s instructions. Flow cytometry was used to characterize the human MSCs. For validation, the following BD Pharmingen™ monoclonal antibodies (mAbs) were used: phycoerythrin (PE) conjugated mouse antibodies with anti-human immunoreactivity for CD34 (#555822), CD44 (#555479), CD45 (#555483), and CD105 (#560839) (BD Biosciences, San Jose, CA). Human MSCs were separately incubated with the above mAbs or mouse IgG isotype control for 30 min at 4°C. Excess mAbs were removed by washing twice with PBS. Cells were resuspended in 0.5 mL PBS to achieve a final density of 2×10^5^ cells prior to acquisition and were then analyzed by a FACSCalibur flow cytometer (BD Biosciences). The surface marker expression of human MSCs thus enriched was assessed by using flow cytometry following 2 passages. The results indicated a high expression of CD44 and CD105 (all > 94% in both MSCs) and a very low expression level of CD34 or CD45 (all < 0.5% in both MSCs) (Figure supplement 1).

### Plasmid construction and transfection

Construction of the pIRES2-Nrf2-DKK1 plasmid, which was used to simultaneously overexpress Nrf2 and DKK1 in MSCs prior to their administration, was based on a pIRES2-EGFP vector (Clontech, Mountain View, CA; #6029-1). cDNA inserts from coding domain sequences (CDS) of the Nrf2 (GenBank accession: NM_006164) and DKK1 genes (GenBank accession: NM_012242) were synthesized by PCR. EGFP gene in the pIRES2-EGFP vector was then replaced by a DKK1 CDS sequence to generate a pIRES2-DKK1 vector. The Nrf2 gene fragment was inserted into the pIRES2-DKK1 vector at the restriction sites of SacI and BamHI. Finally, overexpression (OE) plasmids containing CDS of both human Nrf2 and DKK-1 genes of Nrf2 and DKK1 were obtained by PCR amplification and sub-cloning into an empty pCDNA3.1 plasmid. List of plasmid constructs is detailed in Table supplement 2. Knockdown (KD) of endogenous Nrf2 (#NM-006164-07241504MN), DKK1 (#NM-012242-07241504MN), NQO-1 (#NM-000903-07241504MN), or HO-1 (#NM-002133-07241504MN) expression was achieved by using corresponding human MISSION shRNAs commissioned with Sigma-Aldrich (St Louis, MO). Transfection of plasmids or shRNAs into MSCs was conducted by using a Lipofectamine 3000 system (#1687583; Invitrogen, Carlsbad, CA). Efficiency of genetic OE or KD was verified according to the manufacturers’ instructions.

### Luciferase reporter assay

Approximately 1 kb upstream region of the *Dkk1* gene transcription start site (TSS) containing the wild-type (WT) antioxidant response element (ARE; TGACTCTGC) or mutated ARE (ATCGAGATA) was conjugated to the translation start site in the pGL3-basic vector (Promega, Madison, Wisconsin). HEK-293T cells were plated in 24-well plates 24 h before transfection. Briefly, HEK293T cells reached 70% confluence at the time of transfection. The transfection system is as follows, 450 ng of pcDNA3.1-Nrf2 plasmid, 75 ng of pGL3-basic-DKK1-ARE-WT (or 75 ng of pGL3-basic-DKK1-ARE-Mut plasmid) and 25 ng pRL-TK were co-transfected using 1.5 μg of HG transgene reagent (IBSbio, Shanghai, China). After 48 h of the transfection, cells were harvested and lysed for luciferase assay. Add 20 μl of sample and 20 μl of Firefly Luciferase Assay Reagent to the measurement tube, mix thoroughly and measure the RLU (relative light unit), while setting the cell lysis buffer as a blank control well. Add 20μl of the prepared Renilla Luciferase Assay working solution to the tested sample, and then measure the RLU after thorough mixing. The RLU values detected by Firefly Luciferase are compared with those detected by Renilla Luciferase, and the activation degree of the reporter gene is determined by the ratio.

### Cell culture, reagents and chemicals

All cell culture reagents and consumables were purchased from Gibco (Carlsbad, CA) or Corning Incorporated (Corning, NY). Hydrogen peroxide (H_2_O_2_; #H1009), *N*-acetyl-L-cysteine (NAC; #S0077) and methylthiazolyldiphenyl tetrazolium bromide (MTT; #2128) were products from Sigma-Aldrich (St Louis, MO). Mitochondria-targeted antioxidant (MitoQ) was purchased from MedChemExpress (#HY-100116; Monmouth Junction, NJ). Antibodies for human proliferating cell nuclear antigen (PCNA; #ab18197), Bcl-2 (#ab196495), phosphorylated p38 mitogen-activated protein kinase (MAPK) at Thr180/Tyr182 (#ab4822), total p38 MAPK (#ab31828), phosphorylated p44/42 MAPK (ERK) at Thr202/Tyr204 (#ab214362), total ERK (#ab17942), Dickkopf-1 (DKK1; #ab93017), alpha 1 antitrypsin (αAT; #ab166610), albumin (Alb; #ab106582), Nuclear factor erythroid 2-related factor 2 (NRF2; #ab92946), Cyclin D1 (#ab16663), STAT1 (#ab239360), phosphorylated STAT1 at S727 (#ab109461), STAT3 (#ab68153), phosphorylated STAT3 at Y705 (#ab76315), Heme Oxygenase 1 (HO-1; ab189491), NAD(P)H quinone dehydrogenase 1 (NQO-1; #ab80588) and pan-glyceraldehyde-3-phosphate dehydrogenase (GAPDH; #ab8245) antibodies were provided by Abcam (Cambridge, England). The p38 MAPK inhibitor SB203580 (#S8307) and ERK inhibitor U0126 (#U120) were purchased from Sigma-Aldrich (St Louis, MO). They were added (20 μM) to the cell culture medium 2 h before toxin (e.g. LPS/H_2_O_2_ or ethanol) treatment. Recombinant human tumor necrosis factor-alpha was purchased from PeproTech (Rocky Hill, NJ; #300-01A).

### ACLF mouse model

Male 7-week-old wild-type (WT) C57BL/6J (approximately 21 g) mice were procured from the Guangdong Experimental Animal Center (Guangzhou, China). The ACLF model was established as previously described (Xiang *et al*., 2020). Briefly, mice were injected intraperitoneally (i.p.) with CCl_4_ (0.2 ml/kg twice a week) and then an i.p. injection with *Klebsiella pneumoniae* (*K.P*.) strain 43816 (ATCC, Manassas, VA). All experimental procedures were approved by the Ethical Committee of Shenzhen Third People’s Hospital (SZTPH: 2016-07). For *in vivo* assessment, mouse serum and liver tissue were collected at day 3 after MSC transplantation, since our previous studies showed that on sampling day 3 (an experimentally optimized window for observation) would allow sufficiently informative evaluation on therapeutic effects from drugs or MSCs (Liu et al., 2017; Zeng *et al*., 2015). For *in vivo* viral injection for CKAP4/LRP6 hepatic knockdown, mice were injected via the tail vein with 1×10^12^ genomic copies of AAV8 control or AAV-shRNA (5 per group). Mice were maintained in a 12 h light/12 h dark cycle. After 14 days, mice were fasted for 4 h at the end of the dark cycle and then sacrificed to ensure hepatic downregulation of CKAP4 or LRP6, for comparison with other major organs.

### Cell viability

Changes in MSC viability after specific treatment(s) were measured by using MTT assay. After treatments, cells were washed by sterile phosphate buffer saline (PBS) 3 times and then incubated with 5 mg/mL MTT (Sigma-Aldrich; #M2128) for 4 h, whose reaction products were subsequently dissolved in dimethyl sulfoxide (DMSO, Sigma-Aldrich; #D2650). Absorbance of MTT was measured at 570 nm and pure DMSO was set as a blank control.

### AAV8-shRNA preparation

Adeno-associated virus type 8 (AAV8) was produced by transfection of AAV-293 cells with three plasmids, namely: an AAV vector expressing short hairpin RNA (shRNA) targeting mouse CKAP4/LRP6; an AAV helper plasmid (pAAV Helper); and an AAV Rep/Cap expression plasmid. At 72 h post-transfection, cells were harvested and lysed by following a freeze-thaw procedure. Viral particles were purified by means of an iodixanol step-gradient ultracentrifugation method. Iodixanol was subsequently diluted, and AAV was concentrated by using a 100-kDa molecular-weight cutoff ultrafiltration device. A genomic titer of 2.5×10^12^ - 5×10^12^ infectious units per microliter was determined by real-time quantitative PCR. To construct shRNAs, oligo-nucleotides containing sense and antisense sequences were joint by a hairpin loop followed by a poly (T) termination signal. The sequences targeting mouse CKAP4 (GenBank accession: NM_175451.1) or mouse LRP6 (GenBank accession: NM_008514.4) as used in the experiments were 5’-CCAAGTCTATCAATGACAACA-3’ and 5’-CGCACTACATTAGTTCCAAAT-3’, respectively. The sequence for generating the mock control shRNA was TTCTCCGAACGTGTCACGT. These shRNAs were ligated into an AAV8 vector expressing H1 promoter and EGFP.

### Apoptotic percentage measurement

After treatments, Hoechst 33342 (5 μg/mL, Sigma-Aldrich; #B2261) and propidium iodide (5 μg/mL, Sigma-Aldrich; #P4170) were added simultaneously to each well to stain live MSCs. Cell population was separated into 3 groups: healthy cells only showed a low level of blue fluorescence; apoptotic cells showed a higher level of blue fluorescence, and dead cells showed low-blue and high-red fluorescence. Stained cells were observed and quantified by two independent experimenters without knowing the group assignment. Results were expressed as a percentage of apoptosis (PA): PA = apoptotic cell number/ total cell number × 100%.^1^

### Measurement of oxidative stress in MSCs

CellROX^®^ oxidative stress reagent (Invitrogen, Carlsbad, CA; #C10444) is a novel fluorogenic probe for measuring oxidative stress in living cells. After treatments, CellROX^®^ reagent was added at a final concentration of 5 μM to MSCs, followed by incubation for 30 min at 37°C for fluorescence (green color) measurement by using an inverted fluorescent microscopeIX71 (Olympus microscope, Tokyo, Japan). Positive signals were quantified by ImageJ software (Version 1.52r; NIH, Bethesda, MD).

### Detection of mitochondrial superoxide by flow cytometry

After treatments, MSCs mitochondrial superoxide was measured with 5 μM MitoSOX (Invitrogen; #M36008) for 15 min at 37 °C. Cells were then washed with PBS, treated with trypsin and resuspended in PBS containing 1% (v/v) heat-inactivated FBS. Data were acquired with a FACS Calibur machine (BD Biosciences, San Jose, CA) and were analyzed with the CellQuest analytical software.

### Serum and liver tissue processing and analysis

After animal sacrifice, mouse serum was collected by centrifugation from whole blood samples at 1,000x g for 10 min at 4°C and stored at −80°C. Serum ALT and AST levels were measured by using ALT (SGPT; #A524-150) and AST (SGOT; #A559-150) reagent sets (Teco diagnostics, Anaheim, CA) according to the manufacturer’s instructions. Liver tissue samples were fixed in 10% phosphate-buffered formalin, processed for histology and embedded in paraffin blocks. Hepatic histology and fibrosis were visualized by staining with hematoxylin/eosin (H&E) or Sirius Red using a LEICA Qwin Image Analyzer (Leica Microsystems Ltd., Milton Keynes, UK).

### Western blotting, ELISA, and RT-PCR assays on key hepatic genes

Protein extraction/quantification from MSCs or murine liver tissues, as well as Western blotting assays were conducted as previously described^3^. Parallel blotting of GAPDH was used as an internal loading control. TNF-α protein level was measured by using an ELISA kit from PeproTech (#900-K25) according to the manufacturer’s instructions. Activity changes of Nrf2 (#TFEH-NRF2) were evaluated by using ELISA kits from RayBiotech (Norcross, GA). Human NQO-1 (#ab28947) and HO-1 (#ab133064) protein level changes were quantified by using ELISA kits from Abcam.

### Assay on transplantation safety

To verify the long-term transplantation safety of MSCs in healthy and ACLF mice, we performed a 24-week tumorigenicity study as previously described elsewhere^5^. Healthy 7-week-old C57BL/6J male mice (with or without ACLF induction) received 1 × 10^7^ MRC-5 (negative control; ATCC, Manassas, VA; #CCL-171), 1 × 10^7^ ES-D3 (positive control; #CRL-11632), or 1 × 10^7^ hADMSCs (MSCs group, with or without pIRES-Nrf2-DKK1 plasmid pre-transfection) (12 mice per group for healthy, 18 mice for ACLF group). After 24 weeks or when mice exhibited severe symptoms of dyspnea and minimal activity, mice were sacrificed to assess the extent of tumor formation.

### Statistical Analysis

Data from each group are presented as means ± SD. Unless otherwise stated, statistical comparisons between groups were done by using Kruskal-Wallis test, followed by Dunn’s post hoc test to determine differences in all groups. A value of p < 0.05 or less was considered statistically significant (GraphPad Prism 5.0; San Diego, CA).

## ACKNOWLEDGMENTS

This work was funded by grants from the National Natural Science Foundation of China (82122009, 81970515, 82170605, and 81873573), the Guangdong Natural Science Funds for Distinguished Young Scholar (2019B151502013), and the Guangdong Basic and Applied Research Foundation (No. 2021B1515120069).

## Author Contributions

Hua Wang and Jia Xiao conceptualized the entire study and wrote the first draft of the manuscript. Feng Chen and Zhaodi Che performed, and analyzed most experiments and co-wrote the paper. Pingping Luo, Lu Xiao, Yali Song, Zhiyong Dong, Mianhuan Li, Min Yang, Dongqing Wu and Yi Lv performed *in vitro* and *in vivo* experiments and analyzed the results. Yingxia Liu, Cunchuan Wang, George L. Tipoe and Fei Wang analyzed the results and revised the paper.

## Additional Files

## Supplementary files

Supplementary file 1: Tumor incidence rate after transplantation of hADMSCs in healthy and ACLF mice.

Supplementary file 2: Sequences of Plasmid Constructs

Transparent reporting form

## Data availability

All data supporting the findings of this study are available within the article and its supplementary files. Source data files have been provided for Figures 1 to 6.

## Figure legends

**Figure supplement 1.**
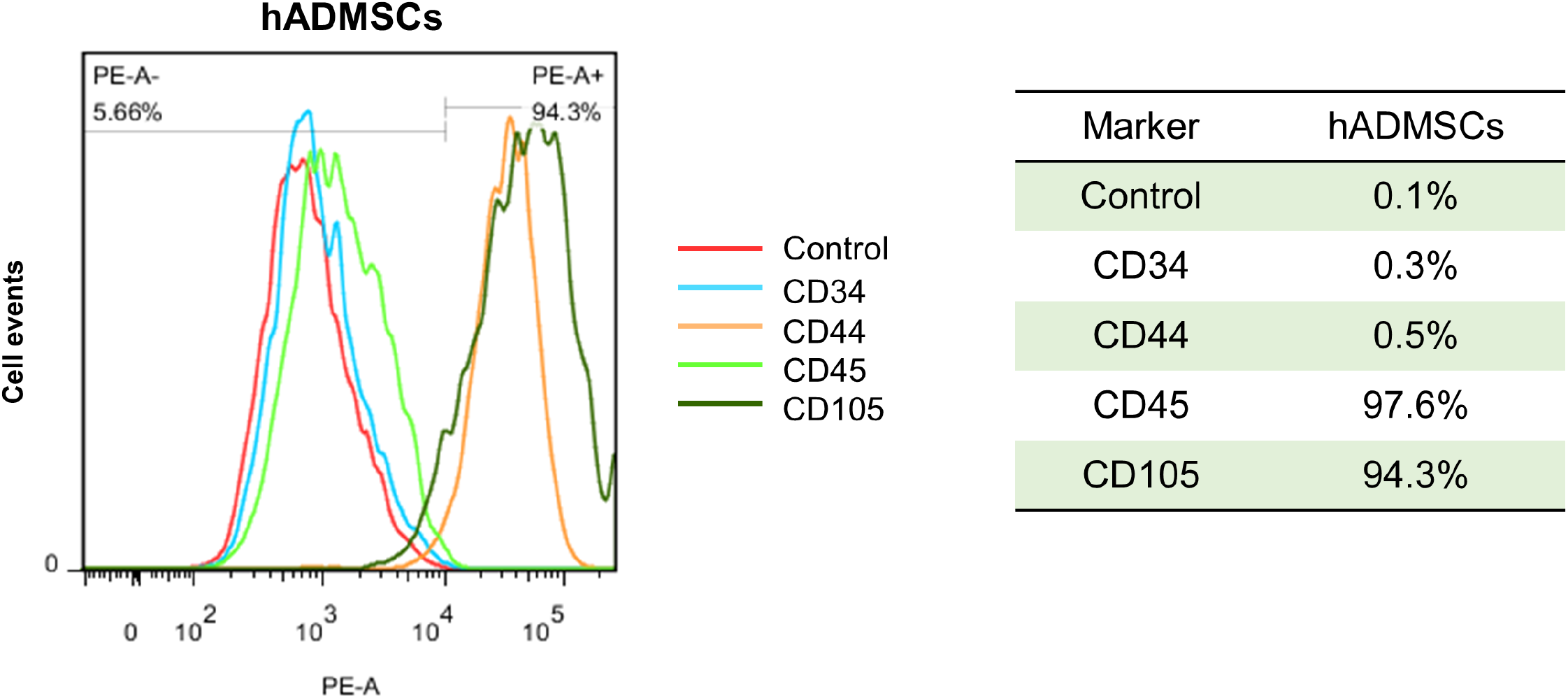
Validation of cell surface markers. Flow cytometry analysis on human adipose-derived mesenchymal stromal cells (hADMSCs) showed their high expression in CD44 and CD105 and low expression in CD34 and CD45.

**Figure 1- figure supplement 2.**
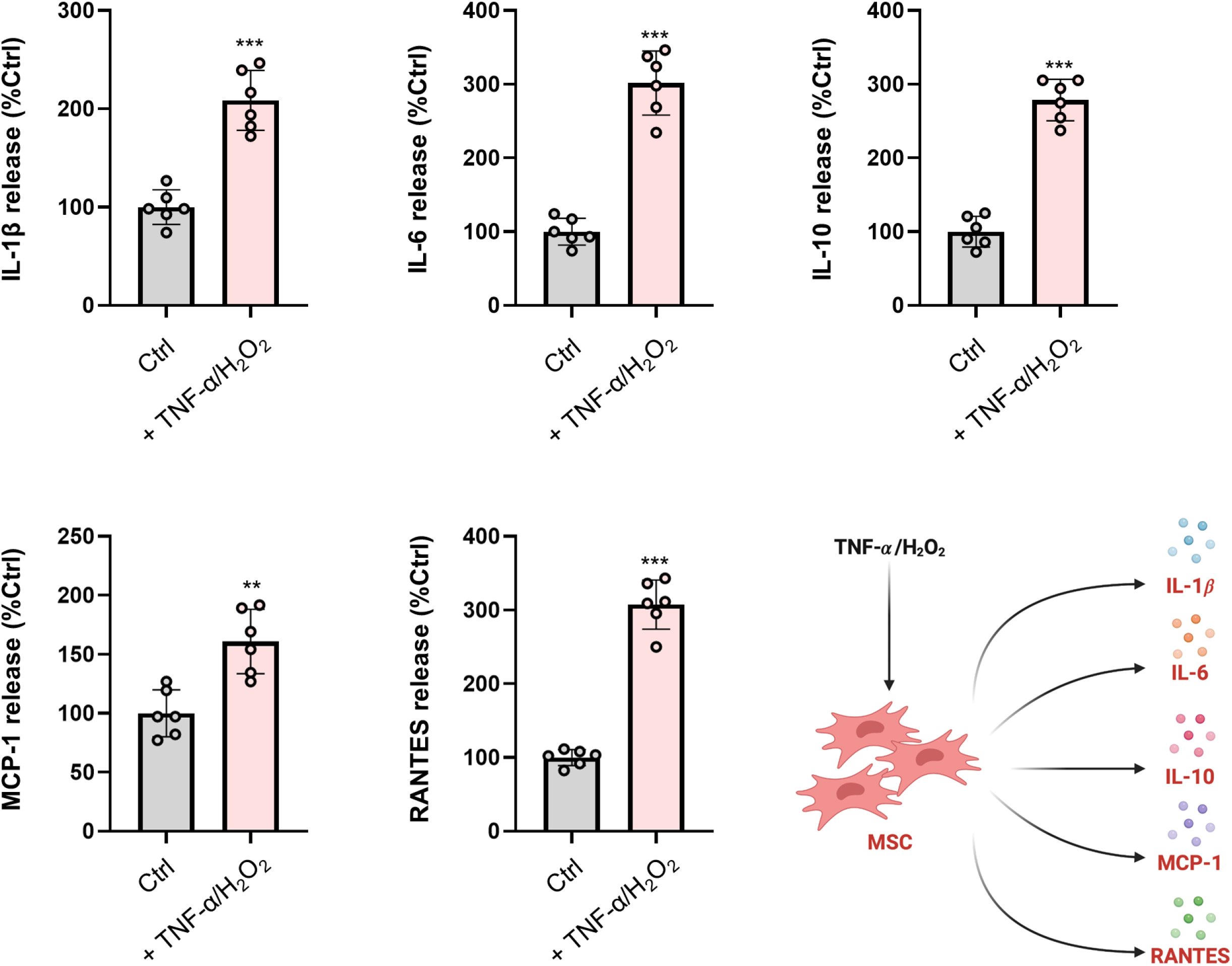
ELISA results of released IL-1β, IL-6, MCP-1,RANTES, and IL-10 protein from MSCs treated with TNF-α/H_2_O_2_ (n = 6). Data are expressed as mean ± SD. *** indicates *p* < 0.001 against the control group.

**Figure 2- figure supplement 3.**
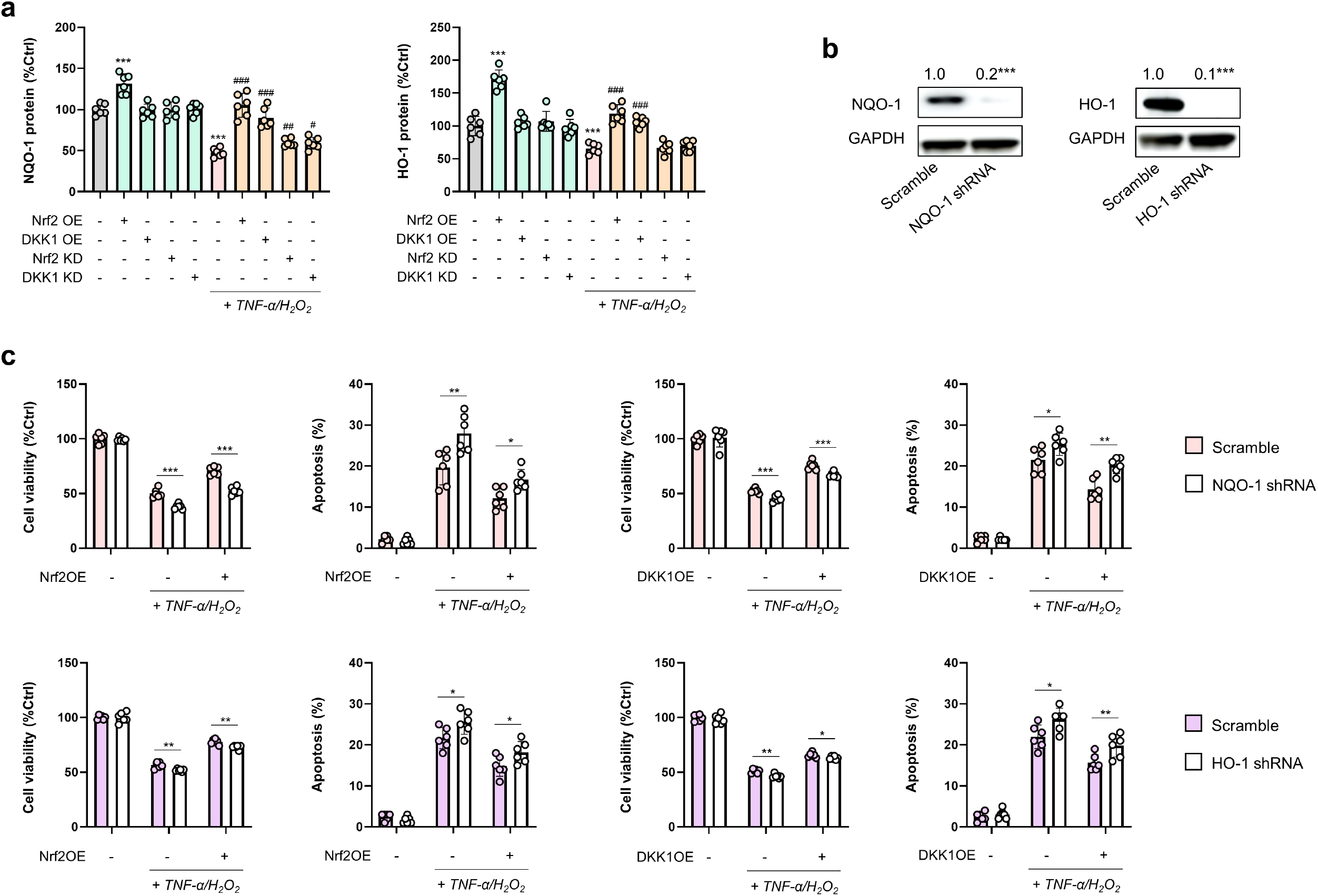
Regulatory loop between Nrf2/DKK1 and NQO-1/HO-1 in MSCs. (A) Changes in NQO-1/HO-1 protein expression of MSCs following TNF−*α*/H_2_O_2_ challenge in the absence or presence of Nrf2/Dkk1 manipulations (*n* = 6). (B) Representative immunoblot results for MSCs when endogenous NQO-1/HO-1 was inhibited specifically by shRNAs. (C) Changes in cell viability or apoptotic ratios of MSCs following TNF−*α*/H_2_O_2_ challenge in the presence or absence of NQO-1/HO-1 inhibition or Nrf2/DKK1 overexpression (OE) (*n* = 6). Data are expressed as mean ± SD. For panels A and B, *** indicates *p* < 0.001 against an untreated MSC group; ^#, ##, ###^ indicate *p* < 0.05, 0.01, 0.001 against a corresponding TNF−*α*/H_2_O_2_ group, respectively. For panel C, *, **, *** represent *p* < 0.05, 0.01, 0.001 between indicated groups, respectively.

**Figure 3- figure supplement 4.**
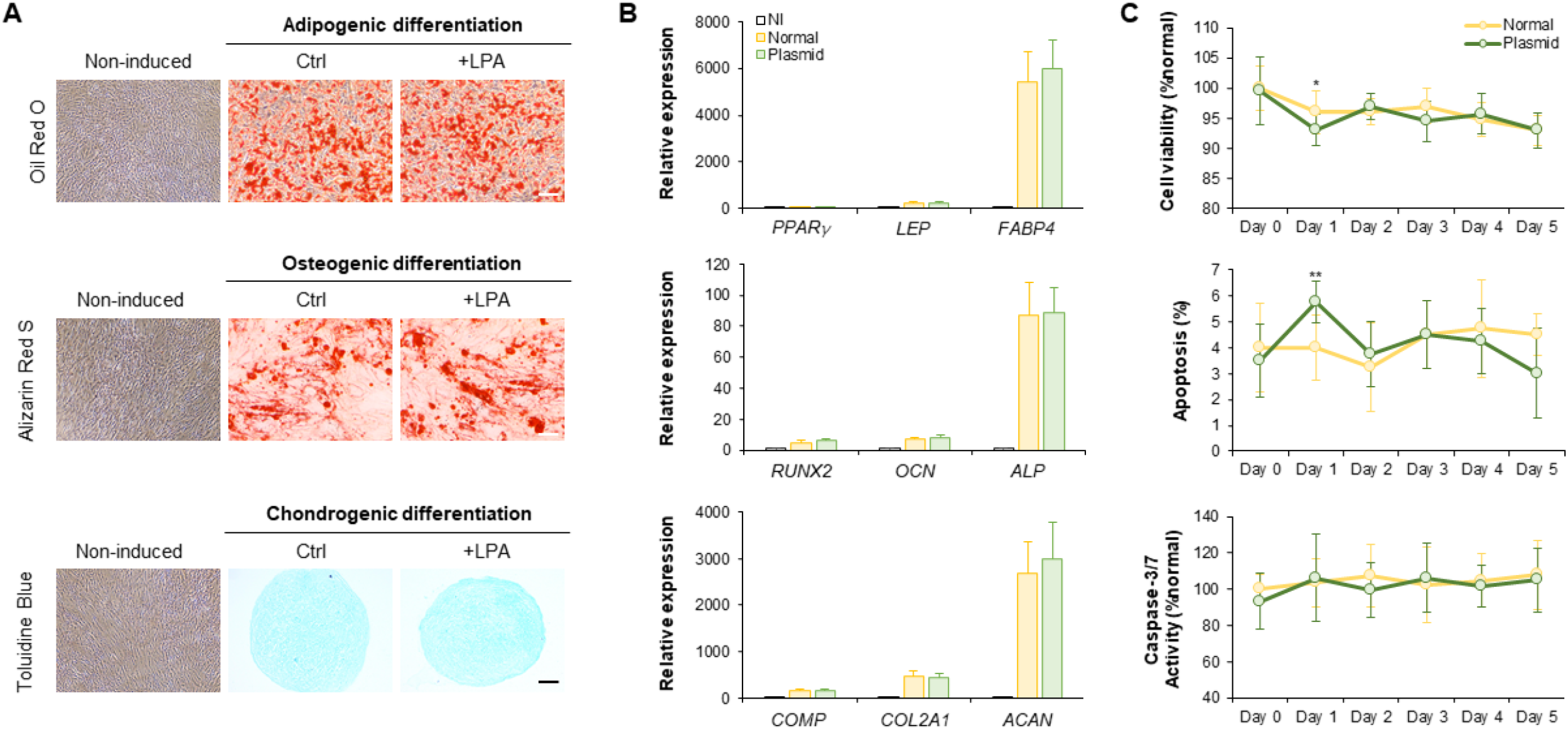
Transfection with a pIRES2-Nrf2-DKK-1 plasmid does not interfere with MSCs transdifferentiating potential or cell status. (A) Representative images of MSCs adipogenic, osteogenic, and chondrogenic differentiation by using commercial standardized protocols. (B) Quantitative RT-PCR measurements of key genes for MSCs adipogenic, osteogenic, and chondrogenic differentiation. (C) Changes in MSCs viability, apoptosis ratios, and caspase-3/7 activities on day 0-5 following transfection with the pIRES2-Nrf2-DKK1 plasmid. All data shown herein are that of MSCs. Data are expressed as mean ± SD. *, ** indicate *p* < 0.05, 0.01 against a corresponding untreated MSCs group, respectively.

**Figure supplement 5.**
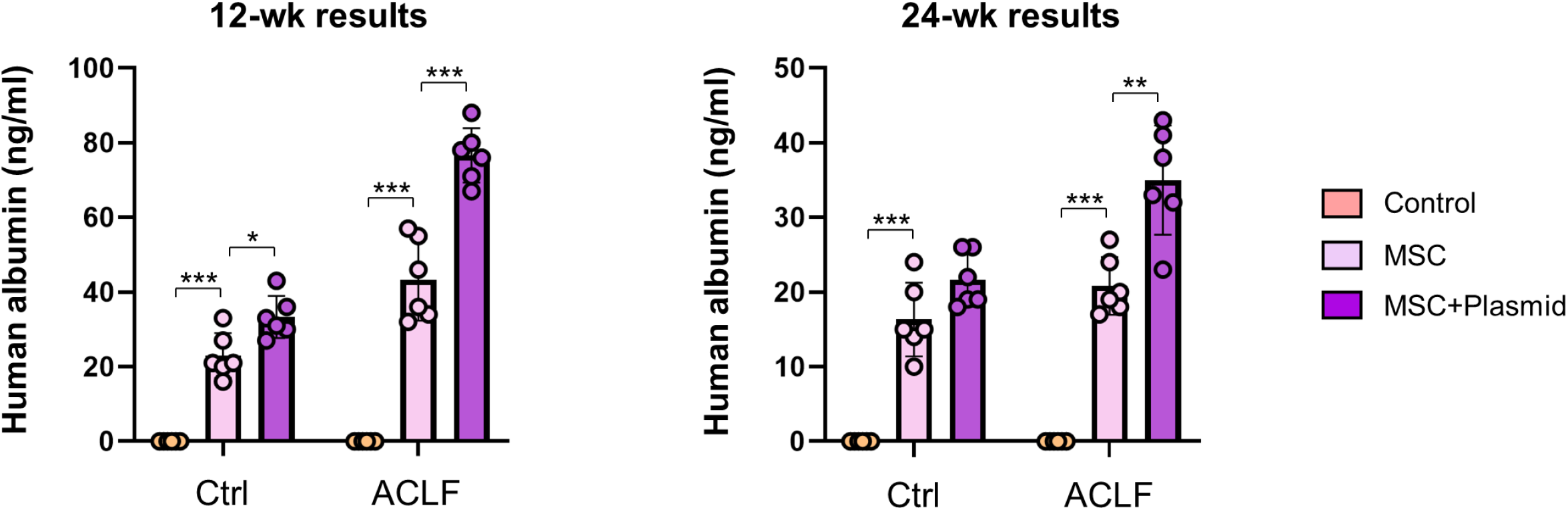
Long-term (12- and 24-week) donor cell function in healthy and ACLF mice transplanted with human MSCs with or without plasmid pre-transfection. Human albumin was determined in the serum by an ELISA assay at weeks 12 and 24 for healthy or ACLF mice (*n* = 6) transplanted with preconditioned MSCs. Mice without MSC transplantation served as negative controls. Data are expressed as mean ± SD. Results are representative of at least 3 independent experiments. *, **, *** represent *p* < 0.05, 0.01, 0.001 between indicated groups.

